# Trade-offs associated with higher winter wheat yields in low soil organic matter cropping systems under climate change

**DOI:** 10.1101/2024.11.30.626142

**Authors:** Jennifer Michel, Vincent Leemans, Markus Weinmann, Iñaki Balanzategui-Guijarro, Jimmy Bin, Simon Biver, Adrien Blum, Rachel Börger, Da Cao, Sok-Lay Him, Gaëlle Kirbas, Jacques Le Gouis, Jordi Moya-Laraño, Mayliss Persyn, Jérome Pierreux, Alice Quenon, Sara Sanchez-Moreno, Sarah Symanczik, Florian Vanden Brande, Dominique Van Der Straeten, Markus Wagner, Matthias Waibel, Anna Xayphrarath, Hervé Vanderschuren, Cécile Thonar, Pierre Delaplace

**Author notes:** Contributed equally (Jennifer Michel, Vincent Leemans, Cécile Thonar, Pierre Delaplace).

## Abstract

Empirical data is key to anticipate the impact of climate change on cropping systems and develop land management strategies that are sustainable while ensuring food security. Here, the combined effects of projected increases in temperature, atmospheric CO_2_-concentrations, solar irradiation and altered precipitation patterns on winter wheat cropping systems were investigated using an Ecotron. Experimental plant-soil systems were subjected to three different climatic conditions representing a gradient of ongoing climate change implementing the weather patterns of the years 2013, 2068, and 2085 respectively. The wheat plants were grown in two differentially manged agricultural soil types: one with long-term low organic matter (OM) inputs and the other one with long-term high OM inputs. In the low OM system, the risk for plant diseases and nitrate leaching was increased, but it outperformed the high OM system with higher yields and lower CO_2_-emissions. Developing high-yielding cropping systems leveraging the CO_2_-fertilisation effect without sacrificing environmental health will therefore require further refined of management practices to improve nutrient cycling and reduce greenhouse gas emissions. One possibility is adapting crop rotations and cover crops to the shorter wheat cycle observed in the future climates to replenish soil nutrients and break disease cycles. Further, in both here studied soil types the wheat plants developed natural coping mechanisms against environmental stressors, such as enhanced root growth and increased levels of proline and silicon. Unravelling the molecular mechanisms that trigger such inherent plant defences is a further interesting target for breeding future crops.

**Graphical abstract:** 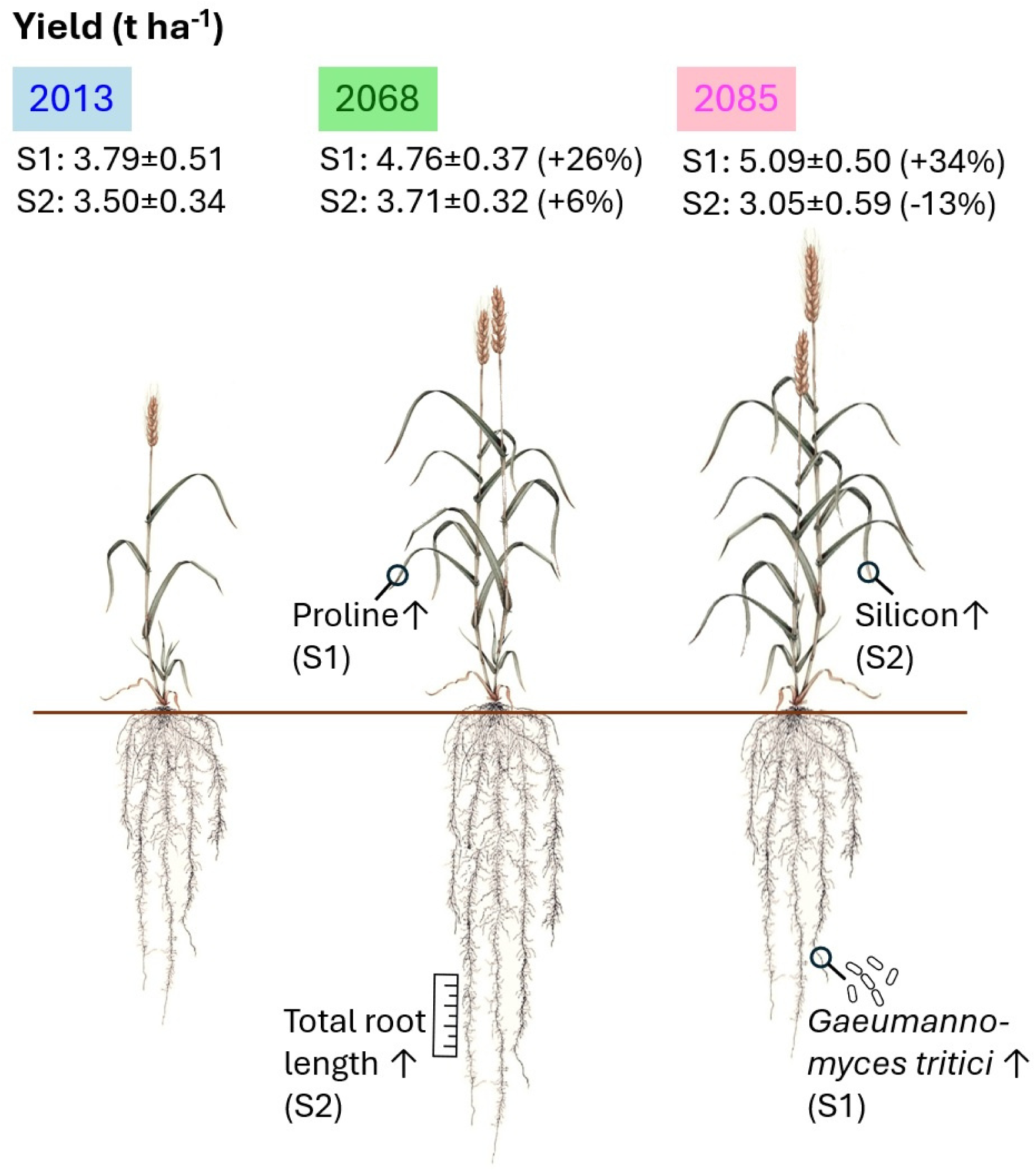

## Introduction

Climate change is expected to significantly impact winter wheat production in Central Europe over the next 100 years, with a theoretical potential for increased yields, but also an increased risk of crop failure following increased heat and water stress (European Environment Agency, 2024). Wheat is currently the most cultivated cereal in Europe, representing 34% of global production, and is the third most cultivated cereal in the world after rice (*Oriza sativa* L.) and maize (*Zea mays* L.), with over 95% of wheat produced worldwide being bread wheat (*Triticum aestivum* L.) (FAO, 2021). While wheat is a staple food in Europe which provides carbohydrates, protein, essential minerals like iron and zinc, and also vitamins like thiamine and pantothenic acid, its large-scale production is critical in terms of its environmental impact, notably nitrate pollution and greenhouse gas emissions (Shewry, 2013; Tilman et al., 2011; Asseng et al., 2019; Cao et al., 2024). Farmers therewith face a triple challenge: increase yields to feed a growing world population, reduce the negative impact of cropping systems on the environment and adapt land management to more and more challenging climatic conditions.

The optimal growth temperature for common winter wheat is 21°C with a non-stressing range between 9 and 22°C, it is sensitive to water logging and drought (Dickin & Wright, 2008; Lamba et al., 2023; Kumar et al., 2023). Under the Representative Concentration Pathways scenario RCP 8.5 W m^-2^ the Western European climate is predicated to be warmer (+3°C), with more (+140 mm) but unequally distributed rain and elevated CO_2_ concentrations (+375 ppm) by the end of this century (IPCC, 2021). While warmer temperatures might initially benefit yields, extreme heat events during critical growth phases (like flowering) can lead to heat stress and reduced yields (Lamba et al., 2023; Riedesel et al., 2023). Rising temperatures, especially combined with elevated atmospheric CO_2_ concentrations are also going to affect growth cycles, potentially leading to earlier wheat maturation and shorter growth cycles (Rezaei et al., 2018). This bears particular risks for winter wheat in central Europe as crops may be exposed to unfavourable weather conditions during their sensitive growth stages, for example heat waves during grain filling. Rising atmospheric carbon dioxide levels as such can have both beneficial and detrimental effects on crops. While increased CO_2_ levels can initially benefit wheat growth through the CO_2_-fertlisation effect, this is only observed under optimal temperature conditions. Beyond these conditions, the benefits diminish, and elevated CO_2_ can decrease yields and protein levels (Myers et al., 2014; Porter et al., 2014). Additionally, elevated CO_2_ levels can promote rhizodeposition, which involves an increased release of root exudates intensifying microbial decomposition of soil organic carbon and potentially increasing CO_2_-emissions from the soil (Jansson and Hofmockel, 2020). Moreover, changes in rainfall, including both increased frequency of heavy rainfall and prolonged droughts, can negatively impact crop growth and ultimately yields, especially if wheats suffer from insufficient water supply in critical growth stages and/or excessive rainfall increases disease pressure and hinders field operations. In general, warmer temperatures and higher humidity levels can facilitate the spread of pests and diseases such as powdery mildew, septoria or take-all, which may lead to poor crop growth or even crop losses if not managed effectively. Similarly, changes in temperature and moisture influence soil microbial communities and nutrient cycling, potentially reducing soil health and fertility over time with negative consequences for the harvested crops.

A further key point to consider is disease propagation in winter wheat cropping systems under climate change, which is influenced by various factors including temperature, humidity, and pathogen biology (Juroszek & von Tiedemann, 2013; Chakraborty & Newton, 2011). Warmer temperatures can accelerate pathogen life cycles and increase disease severity, especially fungal diseases may become more prevalent. Altered precipitation patterns and higher humidity can also promote the growth of diseases such as powdery mildew and take-all (Ghini et al., 2011; Miedaner, 2018). With shifts to the timing of winter and spring, crop phenology and pathogen life cycles may face new combinations with currently unknown consequences for crop vulnerability towards pathogens. Integrated pest management strategies will be crucial for adapting to these changes, but these strategies depend on accurate information about the anticipated disease pressure.

To prevent devastating impacts of climate change on winter wheat cropping systems and anticipate negative consequences it is crucial to have accurate empirical data on future crop growth cycles, plant nutrient requirements and disease propagation. To date, most experiments focus on individual climate change factors such as elevated CO_2_, however one of the cruxes of climate change is the multifactorial aspect of abiotic stresses combining elevated CO_2_, altered precipitation patterns and rising temperatures at varying intensity. Here, an Ecotron facility was used to expose wheat plants growing in two soils with contrasting organic matter management to three climate change scenarios representing the meteorological conditions of the years 2013, 2068 and 2085 respectively. Above and belowground cropping system performance was monitored during the full growth cycle and thus unique insights to the agronomic performance and the environmental impact of the wheat of the future were obtained. The experimental insights provide new information about phenological shifts, disease propagation, plant adaptation strategies, plant nutrient uptake and yield, as well as nitrate leaching and greenhouse gas emissions.

The experiment addresses two main research questions: (i) how do future meteorological conditions impact winter wheat cropping systems and (ii) can low or high organic matter soil management strategies prevent some of the anticipated negative impacts of climate change on crop system performance and yield?

## Methods

### Time travelling with Triticum: Experimental set-up in the TERRA-Ecotron

The experiment was implemented in the TERRA-Ecotron (Gembloux Agro-Bio Tech, University of Liège, Belgium; Roy et al. 2021). The TERRA-Ecotron was built in 2018 and currently has six controlled environment rooms (CERs). The experiment was carried out in mesocosms, which have a soil compartment of 125 L each (cubes of 50x50x50cm), which allows to place nine mesocosms in each CER, resulting in a total of n=54 experimental mesocosms. The first factor “climate” was implemented with three levels, i.e. wheat was grown under the meteorological conditions of the three years 2013, 2068 and 2085 respectively. The second factor crossed with “climate” was “soil management”, which had two levels that is low organic matter content (S1) and high organic matter content (S2). This resulted in a total of six modalities (2013.S1, 2013.S2, 2068.S1, 2068.S2, 2085.S1, 2085.S2). Each modality was implemented with eight replicates, half of which were kept untouched until final harvest for realistic estimation of yield components and the other half was sampled repeatedly during the experiment for destructive measurements such as root growth or leaf elemental composition, each time taking 3-5 plants. In addition, for each soil type in each climate an unplanted control cube was kept to measure baseline soil processes. Having six CERs meant that each climate (2013, 2068, 2085) was replicated in two CERs and the replicate cubes of each of the two soil management type were randomly distributed amongst these CERs (Fig. 1).

**Fig 1.**
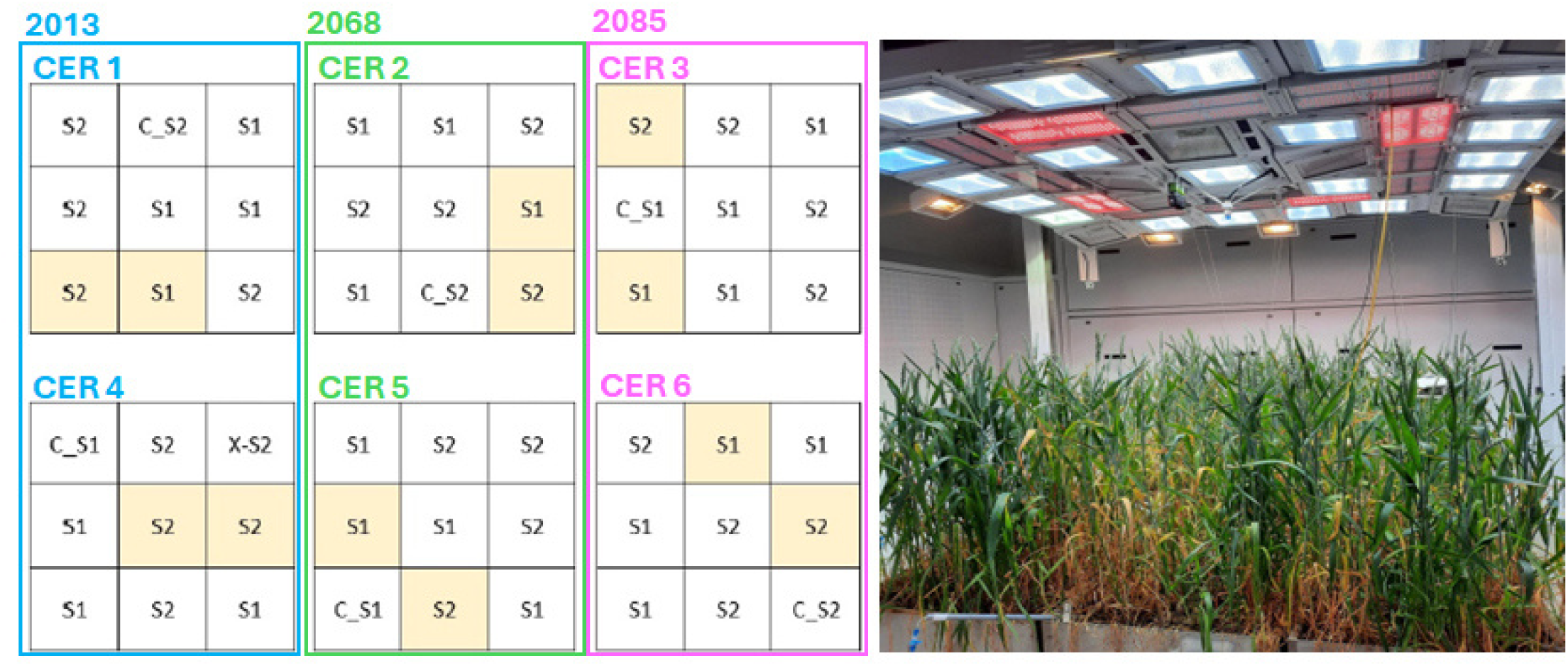
Experimental set-up. Left: Schematic representation of the TERRA-Ecotron which consists of six controlled environment rooms (CERs), in each of which n=9 cubes are placed. Each cube measures 50x50x50cm and contains a soil monolith originating either from the field with low organic matter content (“S1”) or the field with high organic matter content (“S2”). In each CER one soil cube is unplanted (“C_”) while all other cubes are sown with *Triticum aestivum* var. Asory. In each of the CERs one meteorological condition is employed representing either one of the years 2013, 2068 or 2085, with each year being replicated in two CERs (2013: CER1, CER4, 2068: CER2, CER5, 2085: CER3, CER6). Darker shadows of the cubes indicate positioning of a scale underneath the cube. Right: View inside one of the CERs during the experiment: Each CER is equipped with a lightening system combing plasma, halogen and LED lamps which can reach maximum photon fluxes of 1200 µmol m^-2^ s^-1^. Sensors for photosynthetic active radiation (PAR) and irradiance are located at canopy height.

### Three climate scenarios: 2013, 2068 & 2085

Historical data of continuous climate observations from the Ernage meteorological station (50°34’33"N, 4°43’1"E, Belgium, since 1980) was used as well as predicted future meteorological conditions using the Alaro-0 model (Giot et al., 2016). The model ran for the Representative Concentration Pathway (RCP) scenario 8.5 W m^-2^ (IPCC, 2014) and the two time periods 2040-2070 and 2070-2100 (Fig. 2). The three years selected from these predictions (2013, 2068, 2085) align on a continuous gradient of increasing temperature, precipitation, hydrothermal index (HI) and atmospheric CO_2_ concentrations (Table 1). HI is a measure of the relationship between precipitation and temperature, calculated as the ratio of total precipitation to one-tenth of the sum of mean temperatures (Meshcherskaya & Blezhevich 1997). A higher HI indicates wetter and/or cooler conditions, suggesting more favourable moisture availability relative to temperature, which can benefit crops and vegetation. Conversely, a lower HI signifies drier and/or warmer conditions, which may indicate drought stress or reduced water availability. The Ecotron reproduces the simulated weather conditions at a very high temporal resolution, where the key environmental parameters such as sun light intensity and temperature are adjusted every five minutes. By these means, diurnal and seasonal variabilities are accurately reproduced, and plant behaviour can be studied under realistic climate scenarios. For example, the irradiation includes natural sun rise and sun set patterns. The main climatic components that were manipulated for each year in this experiment were atmospheric CO_2_ concentration, temperature and precipitation. CO_2_ concentrations were along a gradient for the three years, with 420 ppm in 2013, 550 ppm in 2068 and 775 ppm in 2085. The historic reference year 2013 was characterised by a mean temperature of 7.59°C during the wheat growth cycle, with a mean precipitation of 2.12 mm d^-1^ (HI=3.99). Interestingly, this year is the most extreme in terms of maximal and minimal temperature (26°C, -7.6°C), and also the year with the longest periods outside the optimum range for wheat in terms of cold and hot days (n=94 days below 4°C and n=9 days above 22°C). The year had a long cold winter from approximately 51-176 days after sowing, during which the soil was frozen and therewith the water present in the soil was not readily available to plants. The year 2013 also had the largest number of rain-free days (n=135), but the lowest number of rainy days (n=57 days with precipitation >=4mm). The 2068 climate was characterised by mean temperature higher than 2013 with 10.17°C but still had a significant winter period with n=40 days where temperatures dropped below 4°C. Mean precipitation was slightly higher than in 2013 with 2.4 mm d^-1^ and only n=4 days without rain, but more rainy days (n=62). The climate of the year 2085 was the smoothest in regards of temperatures around 12.1°C and very rarely below 4°C or above 22°C. The winter period was short with only n=3 days below 4°C. In regards of precipitation, 2085 was the wettest scenario with n=72 days of rain (>=4mm). Interestingly, all three years experience at least one day with very high rain (>30mm), though these maximum rain events occurred at different moments during the season: in March for 2068, in May for 2085 and in July for 2013.

**Fig 2.**
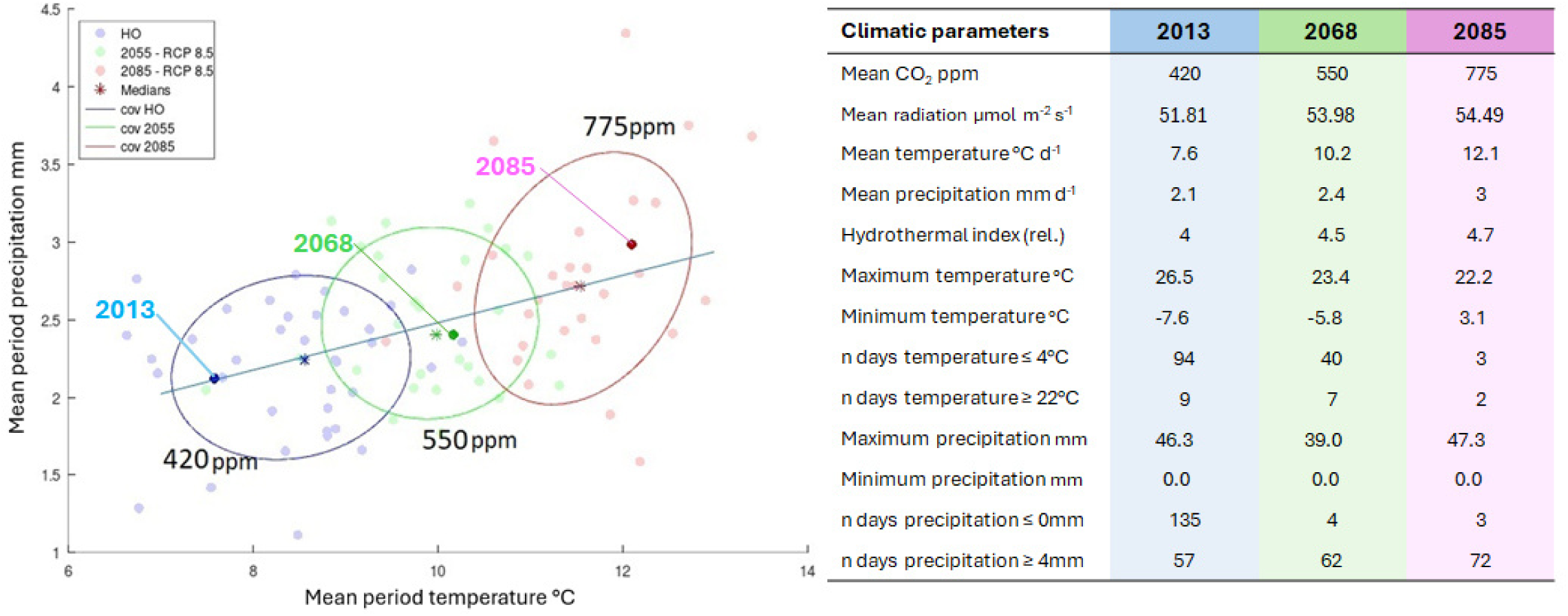
Climate scenarios. Left: Wheat agroecosystems were exposed to the meteorological conditions of three years representing the present climate (2013), the near future (2068) and a farther future (2085). The three years align on a continuous gradient of hydrothermal index with increasing temperature, precipitation and atmospheric CO_2_-concentrations. The climate scenarios are based on continuous meteorological observations from the Ernage weather station (50°34’33"N, 4°43’1"E, Belgium, since 1980) and the predicted future climates were simulated using the Alaro-0 model. The historic observations (HO) cover the period 1981-2017 (blue) and the model ran for the Representative Concentration Pathway (RCP) scenario 8.5 W m^-2^ for the two time periods 2041-2070 (green) and 2070-2100 respectively (rosé). Each dot represents a year and the ellipses represent the 95% confidence interval of the three periods. Right: Table summarizing the key climatic parameter of the selected years.

**Table 1.**
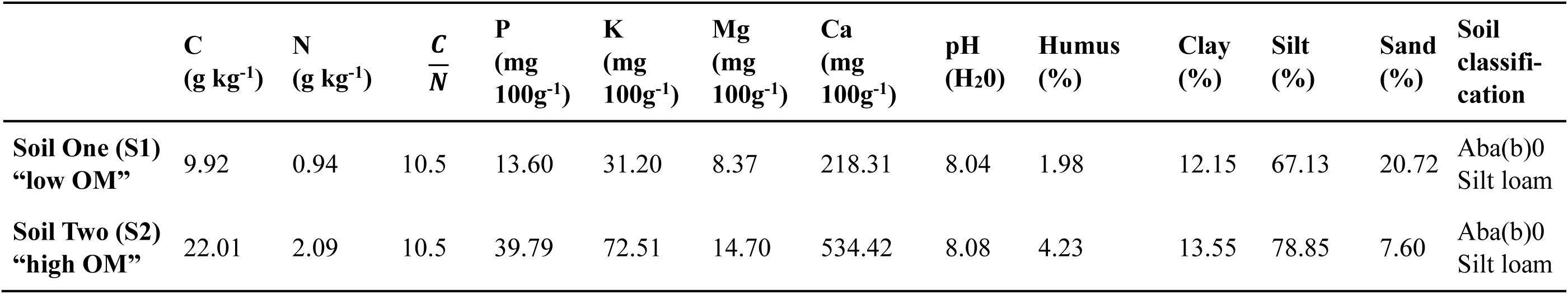
Physicochemical characterisation of the two soil types at the beginning of the experiment, one composite sample per soil type, sampling depth: 0-20cm. Abbreviations: OM: organic matter, C: carbon, N: nitrogen, P: phosphorous, Mg: magnesium, Ca: calcium.

### Two soil types with different organic matter content

The experiment tests two soils which are very closely related in terms of pedogenesis, both originating from the Walloon Brabant in Belgium (S1: 0°38’35.1474”N, 4°37’22.0123”E, S2: 50°39’12.8668”N, 4°38’10.7664”E). In this area limestone formations are prevalent and soils are often clay-rich, which helps retain moisture and nutrients making them very suitable for agriculture (Gentile et al., 2009). Both soils are classified as Aba(b)0 in the Belgian soil classification system and characterised as silty loam (Cigale, 2021). Both soils are from fields that are under agricultural management since generations and implement the regular local crop rotations with winter wheat and root vegetables mainly. In recent years, cover crops with plants such as phacelia, oats, radish and clover have always been grown between two main crops in S2. In both fields standard and reduced tillage was applied regularly, as well as commercial fertilisers, green and brown manure and occasionally herbicides (supplementary material SM1). The main differences between the two soils is that soil two (S2) has received significantly higher quantities of organic matter than soil one (S1). While the soils have similar C:N of 10.5 and pH just above 8, these higher organic matter inputs left S2 with more than doubled humus, carbon (C), nitrogen (N), phosphorous (P), potassium (K), magnesium (Mg) and calcium (Ca) contents as compared to S1 (Table 2). Another difference is in the soil texture, which is sandier for the low organic matter content soil S1. In each field, a total of n=27 cubes were sampled in November 2022 and moved to the Ecotron. Soils were sampled as undisturbed soil monoliths (50x50x50cm) with a surface of 0.25 m^2^ and each weighing approximately 200 kg. Monoliths were taken to realistically represent field conditions in the Ecotron, avoid disturbance of sensitive soil organisms and keep the soil structure intact. One cube of each soil type in each CER was placed on a scale to monitor the weight of the cubes and improve estimates of evapotranspiration.

**Table 2.**
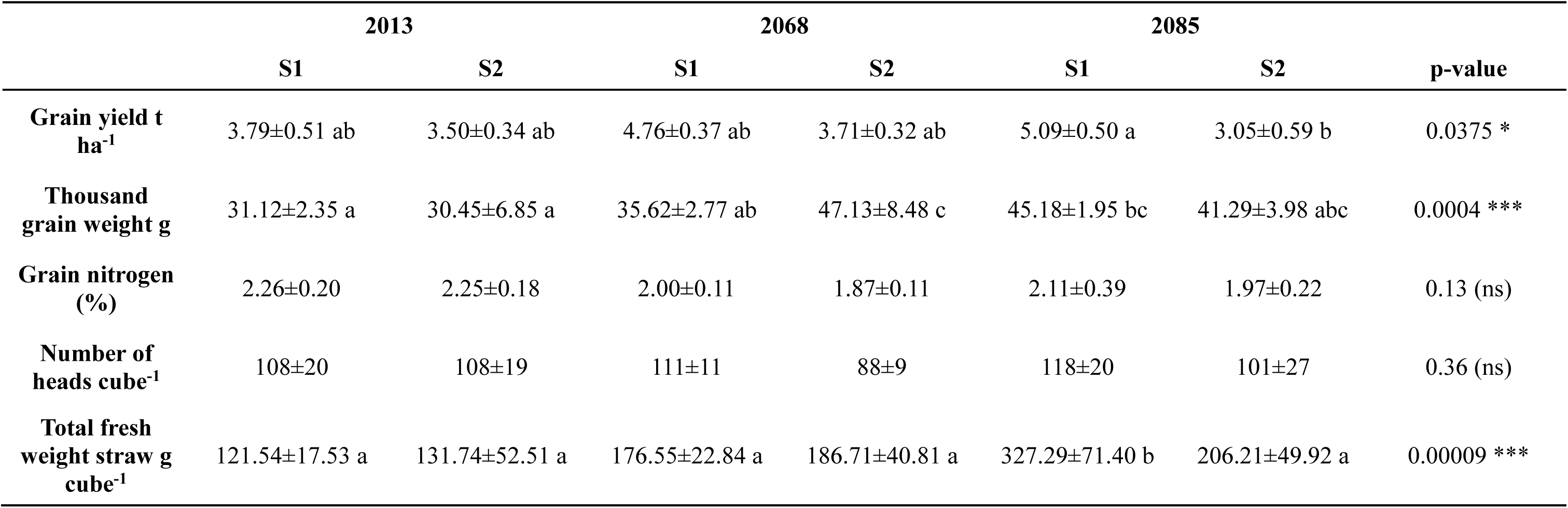
Yield components for winter wheat (*Triticum aestivum* var. Asory) grown in two differentially managed soil types (low OM content S1 and high OM content S2) in the meteorological conditions of the years 2013, 2068 and 2085. Probability values following analysis of variance with asterisks indicating significance levels at <0.001***, ≤0.05*, >0.05 not significant (ns) and letters indicating grouping based on TukeyHSD test.

### Crop management and monitoring

During an initial acclimatisation period of three weeks, the soil cubes were kept under respective climates in the CERs of the Ecotron. At the time of the monolithic sampling for the Ecotron trial the S2 field was pre-cropped with a radish mix, in grass cover with a dense layer of mulch in the sub-surface (wheat straw). There was no cover crop sown in the S1 field prior to sampling the cubes, but a straw layer had also been incorporated to the soil before the sampling. The soils in all cubes were weeded and surface tillage (raking to 15cm depth) was applied during the acclimatisation period. On 23/12/22, which corresponds to 01/11 Ecotron date, 54 cubes were planted with winter wheat (*Triticum aestivum* (L.) var. Asory) at a density of 308 seeds m^-2^ (77 seeds per cube) as recommended for this region (Livre Blanc Céréales, 2022). Cubes were weeded manually when needed and no herbicides were applied. At the beginning of the experiment, metaldehyde pellets were applied to control molluscs. The pellets were placed in the reversed lids of 50 ml Falcon tubes which were then placed on the soil to minimise impact on soil chemistry and other soil organisms (Birkhofer et al., 2008; Iglesias et al., 2003). After germination, plant development was closely monitored and several agronomic and environmental parameters were regularly recorded (Fig. 3, supplementary material SM4). For the aboveground compartment, plant BBCH growth stages, plant height and leaf area index (LAI) were measured to quantify plant growth and maximum quantum efficiency of photosystem II (Fv/Fm) was measured as an indicator of plant performance (Meier, 2001; Vlaović et al., 2020). At three time points (BBCH30/50/80) foliar silicon and foliar proline were determined as important molecules in plant stress response (Hayat et al., 2012; Wang et al., 2017). For the belowground compartment, soil microbial biomass and total root length were measured at the same time points and root infestation with the fungus *Gaeumannomyces tritici* (take-all disease) was quantified once at 230 DAS (Campbell et al., 2003; Seethepalli et al.,2021; BBA, 1999). Unless prevented by drought conditions, interstitial soil pore water was extracted weekly to quantify freely available nitrate in aqueous soil solution and glucose equivalents as indicators of root exudation (Folegatti et al., 2005; Yemm & Williams, 1954). During the experiment, both soils in all climates were fertilised with ammonium nitrate (N: 27%), which was applied in three doses according to plant growth stage, namely at the end of tillering/stem-elongation, 2. node and flag leaf in each climate respectively. The quantities of N-fertiliser varied between 150 and 205 kg N ha^-1^ for the different soils and climates as a function of the mineral N measured in the different soils at the end of winter, with S1 receiving approximately 50 kg N ha^-1^ more than S2 (supplementary material SM3). Plants were harvested when fully ripe (BBCH89) and for each cube aboveground biomass was determined (straw/leaves/heads), as well as number of heads, grain fresh weight, grain humidity, grain yield at 14% moisture, thousand grain weight and grain nitrogen content.

**Fig 3.**
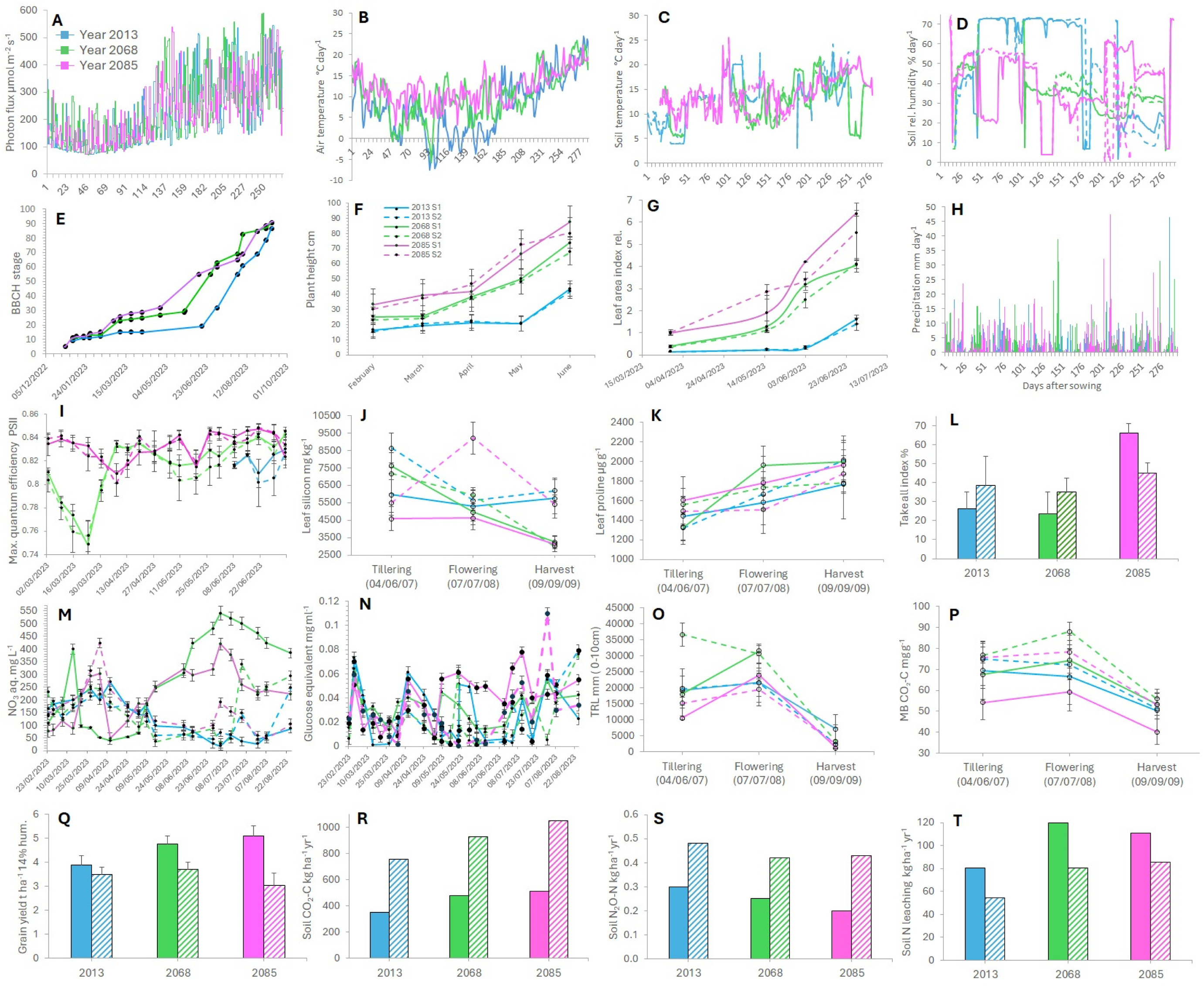
Environmental, crop and soil parameters of two management systems (S1: continuous line/fill and S2: dotted line/hatched area) in the meteorological conditions of the years 2013 (blue), 2068 (green) and 2085 (rosé). Details of each measurement method and replication for each parameter in SM4. A: Photon flux µmol m^-2^ s^-1^, B: Air temperature °C day^-1^,C: Soil temperature °C day^-1^, D: Soil relative humidity % day^-1^, E: BBCH scale plant growth stages, F: Plant height cm, G: Leaf area index (LAI), H: Precipitation mm day^-1^, I: Maximum quantum efficiency of photosystem II (Fv/Fm), J: Leaf silicon mg kg^-1^, K: Leaf proline µg g^-1^, L: Take-all index (% *Gaeumannomyces tritici* infestation), M: Nitrate (NO_3_) mg L^-1^in aqueous soil solution, N: Glucose equivalent in aqueous soil solution mg ml^-1^, O: Total root length (TRL) mm for 0-10cm soil depth, P: Microbial biomass (MB) CO_2_-C mg g^-1^, Q: Grain yield t ha^-1^ at 14% grain humidity, R: Annual soil CO_2_-C budget kg ha^-1^ yr^-1^, S: Annual soil N_2_O-N budget kg ha^-1^ yr^-1^, Annual soil N leaching kg ha^-1^ yr^-1^, A-D, H: Daily measurements, E: BBCH growth stages evaluated for all cubes and only one value noted per modality, F-G, I-Q: Dots represent means ± SD of n=4 or n=8, R-T: Data modelled with DNDC and only one value per modality.

### DNDC model

DNDC (i.e. DeNitrification-DeComposition) is a process-based computer simulation model of carbon and nitrogen biogeochemistry in agro-ecosystems (Giltrap et al., 2010; Gilhespy et al., 2014). DNDC predicts soil environmental factors, C sequestration, and emissions of C and N gases primarily based on microbe-mediated biogeochemical processes, including decomposition, nitrification, denitrification, fermentation, and methanogenesis (Li et al., 1992a, 1992b, Li, 2000). DNDC simulates these processes based on the activity of different functional groups of microbes under different environmental conditions including temperature, moisture, pH, redox potential (Eh) and substrate concentration gradients in soils. For example, nitrification is modelled as first-order process based on soil ammonium concentration (NH_4_^+^) under aerobic conditions and nitrous oxide production (N_2_O) is modelled as a fraction of the overall nitrification rate. Soil Eh is calculated with the Nernst equation at a daily time step following soil saturation and then used to determine anaerobic microbial group activity under a given set of soil conditions. Anaerobic microbial group activity is then modelled using standard Michaelis-Menten-type kinetics. The DNDC model has been extensively evaluated against datasets of trace gases fluxes that were measured worldwide (Gilhespy et al., 2014; Giltrap et al., 2010). To access the accuracy of the model for this experiment, the model’s predicted outputs were compared with measured Ecotron variables and the coefficient of determination (R²) was used as a measure of goodness of fit (supplementary material SM5).

### Statistical analysis

The effects of climate and soil type on the empirically measured parameters (supplementary material SM4, lines 1-12) were assessed using linear mixed-effects models (LMM) with climate and soil as fixed effects, and random intercepts for time and CER to account for the within-subject correlation of repeated measures and replicate rooms. Post-hoc comparisons using estimated marginal means (EMMs) were used where appropriate to elucidate differences between the levels of each fixed effect, i.e. the interaction between climate and soil (2013/2068/2085xS1/S2). For yield components (Table 3, Fig. 3Q) and take-all index (Fig. 3L) analysis of variance (ANOVA) and post-hoc Tukey’s HSD were used to identify differences between climate and soil type. The parameters derived from the DNDC model, that is upscaled budgets of CO_2_, N_2_O, N leaching and transpiration, have only one observation per modality (Fig. 3R-T). They are descriptive but were indicatively compared via Krustkal-Wallis test for climate and soil individually. To reduce the overall dimensionality of the dataset to better understand what the core characteristics of each cropping system in each climate are, probabilistic principal component analysis (pPCA) was performed (Fig. 4). pPCA extends traditional PCA by incorporating a probabilistic framework, which allows the estimation of principal components to identify underlying latent structures in the presence of noise (Tipping & Bishop, 1999). Statistical analysis was carried out using R 4.4.1 (R Core Team, 2024) with the additional packages car (Fox & Weisberg, 2019), emmeans (Lenth, 2024), ggplot2 (Wickham, 2016), lmerTest (Kuznetsova et al., 2017), missForest (Stekhoven & Buehlmann, 2012) and multcompView (Graves et al., 2024).

**Fig 4.**
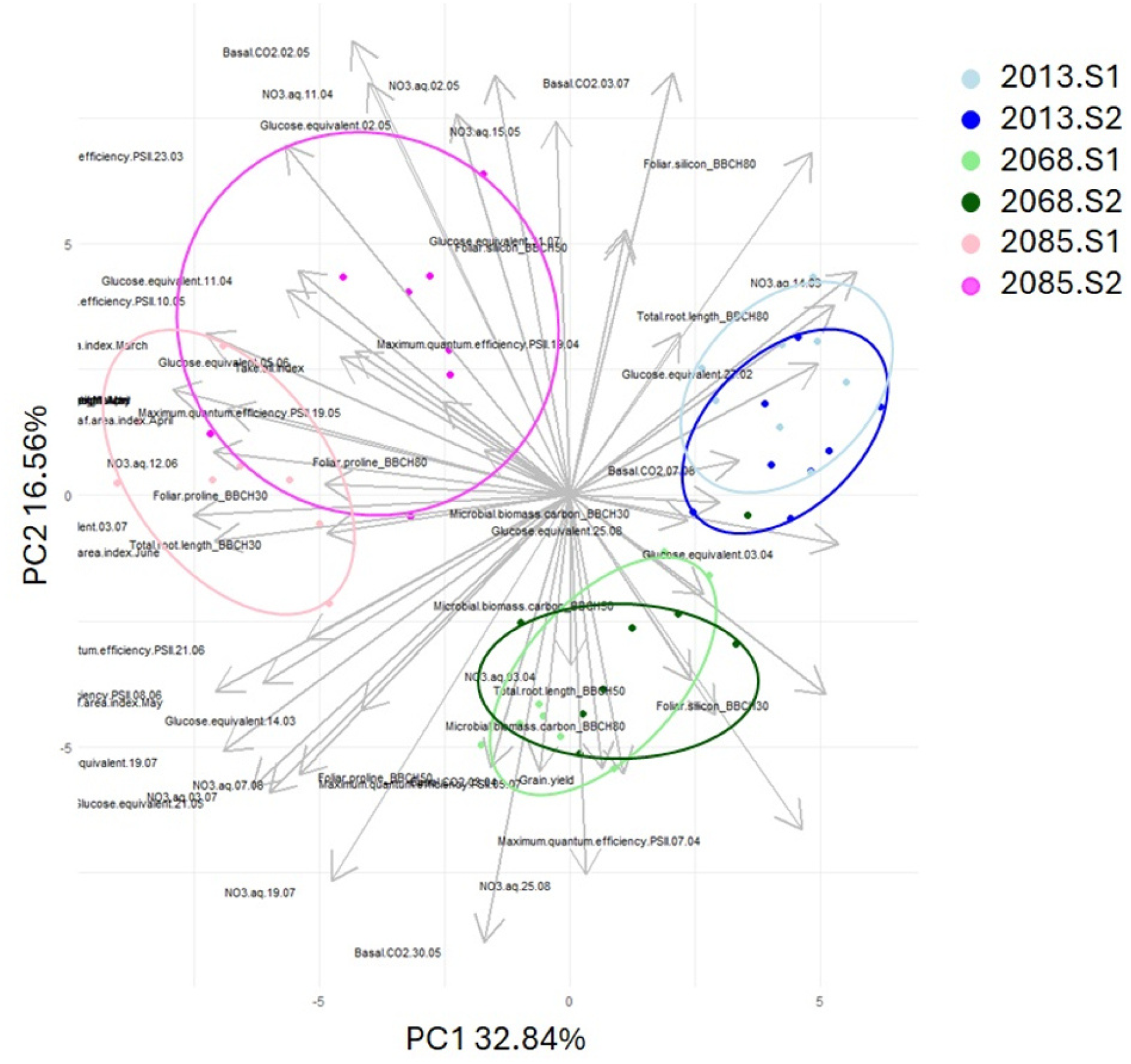
Ordination of agronomic and environmental parameter in probabilistic principle component analysis (pPCA) and clustering by simulated year and soil type. Main parameters which distinguish the groups: 2013.S1: glucose equivalent, take-all index; 2013.S2: leaf area index; 2068.S1: foliar proline, microbial biomass carbon, total root length; 2068.S2: basal CO_2_, maximum quantum efficiency PSII; 2085.S1: plant height, aq. NO_3_, total root length; 2085.S2: foliar silicon, plant height (please see supplementary material SM4 for further details on vector loadings).

## Results

### Phenological advance in future meteorological conditions

Warmer temperatures and more intense rain, but overall less stressful weather with higher atmospheric CO_2_ concentrations meant that crops grew faster and bigger the further in the future the climate scenario was (Fig.3E-G). The phenological advance manifested from February onwards, with plants maturing faster in 2068 and 2085 (BBCH p<0.0001). There was a significant effect of soil type in interaction with climate for plant height (p<0.0001), with taller plants the further in the future the climate scenario. Notably, plants in 2085 were almost twice as high as in 2013 (Fig.3F) and plant height was one of the main characteristics of the 2085 cropping systems on both soil types in the ordination (Fig. 4, supplementary material SM6). In 2013, plants in S2 grew taller than plants in S1 (2013.S1: 19.3±0.445, 2013.S2: 20.8±0.476). The soil type effect was reversed for the two future climates where plants grew taller in S1 compared to S2 and (2068.S1: 25.6±0.445, 2068.S2: 24.0±0.445, 2085.S1: 39.4±0.445, 2085.S2: 37.2±0.445). Small leaf area index (LAI) was one of the main characteristics of the 2013 cropping systems, particularly S2 (Fig. 3G, Fig. 4, supplementary material SM6) and considerably increased in the future climates (p<0.0001).

### Early harvest and yield components

The phenological advance entailed earlier harvests in the future years, with approximately two weeks between each of the climate scenarios. Plants in 2068 were harvested first in early July, followed by 2085 in late July, while 2013 was harvested beginning of August (all dates refer to Ecotron months). Visually distinguishable differences in plant developmental stages between the two soil types were minor and plants from both soils were harvested simultaneously once all grains were fully ripened for the respective year.

Grain yield was always higher in S1 as compared to S2 (Fig.3Q, Table 3), with an overall significant soil type effect and minor differences between climate scenarios (p=0.04). Most notably, yield increased proportional to the increasing hydrothermal index across the three years for S1, meaning that for this soil type the further in the future the climate scenario, the higher the yield. The globally highest aboveground biomass and also highest grain yield were achieved for S1 in 2085, while the lowest grain yield was recorded for S2 in 2085. Thousand grain weight was also significantly affected by soil and climate (p<0.0001) with lowest values in 2013 and highest values for S2 in 2068 (Table 3). Grain nitrogen was not significantly different between soils or climates, however trended to be higher in S1 as compared to S2 and was highest in 2013 (p=0.13).

### Plant health

The maximum quantum efficiency of photosystem II, represented as Fv/Fm (Fig.3I), is a crucial parameter in plant physiology with values around 0.8 considered optimal and lower values indicating stress leading to impaired PSII efficiency (Maxwell & Johnson, 2000; Baker, 2008). Fv/Fm is measured at leaf level and requires significantly wide leaf area, which is why the measurement was initially not possible in 2013. For the data available, Fv/Fm was not significantly different between soil types, but significantly different between climates (p<0.002), with on average higher Fv/Fm in 2085 than in 2068 (than in 2013 when measured). A notable drop in Fv/Fm was recorded for 2068 compared to 2085 at the beginning of the experiment (March Ecotron month) and overall lowest Fv/Fm was measured for S2.2068 which distinguished this cropping system (Fig. 4, supplementary material SM6).

Leaf silicon and proline levels as key indicators of plant stress and adaptation mechanisms were also characterised (Fig. 3J, 3K). There was notably elevated foliar silicon in S2.2085 at BBCH50 and elevated proline in S1.2068 at BBCH80 (Fig. 4, supplementary material SM6). Overall, leaf silicon levels were most distinguished 2085 (p=0.02) with the lowest levels for S1.2085 and the highest foliar silicon for S2.2085 (Fig.3J). Foliar silicon tended to be globally higher in S2 than in S1 (p=0.095). Foliar proline also stood out in 2085 (p=0.07), with an opposite trend to silicon, having lowest values in S2.2085 and highest in S1.2085 together with S1.2068 (Fig.3K).

Take-all disease is a common struggle in winter wheat plantations (Palma-Guerrero et al., 2021) and was quantified at BBCH80 in this experiment (Fig.3L). Symptoms of root infestation with the fungus *Gaeumannomyces tritici* (take-all index) were increased in 2085, especially in S1, but overall no significant climate or soil effect was detected (p=0.06).

### Rhizosphere processes and environmental impact

Decreased total root length (TRL) was amongst the main characteristics of the two future cropping systems on S1 in the ordination (Fig. 3O, Fig. 4, supplementary material SM6). TRL differed particularly at the early stages of plant development where it was higher in S2 compared to S1, but TRL was in frequentist approach not significantly different (p=0.08). Microbial biomass in the root zone was quantified at three time points corresponding to BBCH 30/50/80 (Fig.3E), where it was always higher in S2 compared to S1 (p=0.03). Microbial biomass was one of the main characteristics of the S1.2068 cropping system, together with glucose equivalent (Fig. 4, supplementary material SM6). Detectable glucose levels averaged around 0.038 mg ml-1 in all modalities and were one of the main characteristics of the S1.2013 cropping system (Fig.4, supplementary material SM6). There was a periodical variation in glucose concentrations with a common low level during winter and then varying peaks according to climate and soil type (Fig. 3N). The overall lowest levels of glucose were detected in S1.2013. Later in the experiment, the largest spikes were detected in the 2085 climate, with the all-time highest peak at the beginning of August in S2.2085 (Fig.3N). Freely available nitrate presented high temporal variation just as did glucose levels (Fig.3M), yet nitrate in soil solution was overall significantly higher in the two future climates (p=0.02) and globally not significantly different between soil types. Most notable spikes in free nitrate were detected in S1.2068 and S1.2085 (Fig. 3M) and according to the DNDC model the risk of nitrate leaching was always higher in S1 compared to S2 (Fig.3T), but not statistically significant for soil nor climate. The DNDC model also predicted both higher soil CO_2_ emissions and higher soil N_2_O emissions for S2 compared to S1 (both p=0.05). Comparing predicted GHG emissions across the three climate scenarios revealed an overall trend towards decreasing soil N_2_O and increasing soil CO_2_ emissions with increasing hydrothermal index (Fig.3R,3S).

## Discussion

### Yield and soil organic matter: Less is more?

In all three climate scenarios, plants in the low organic matter content soil S1 yielded more grains than plants in the high organic matter content soil S2 (p=0.007). While the S1 soil received ¼ more mineral N fertiliser than S2, the S2 soil was characterised by 2x higher total C and N contents than S1 (Table 2, supplementary material SM3). The proportional increase in yield with the hydrothermal index for S1 supports previous findings that optimal moisture and temperature conditions can lead to substantial yield improvements in the future (Wilcox & Makowski, 2014). That this effect was observed with S1 but not with S2 aligns with other research showing that soil properties play a key role in wheat productivity, especially as climate conditions fluctuate (Zhao et al., 2022). However, the better realisation of the yield potential under future climates in S1 adds some important nuance as to whether increasing organic matter and soil organic carbon per se is an advisable farming practice. It may be that in soils with higher organic carbon content and larger microbial communities, the competition between soil microbes and plants is enhanced to a degree which prevents optimal plant nutrient uptake and growth (Kuzyakov & Xu, 2013). For harnessing the yield potential in this study, a tipping point was identified for S2 where yields increased for the near future, but decreased strongly for the far future scenario (Fig.3Q, Tabel 3). The result of this study with a yield gap between the two soil types aligns with a mesocosm experiment in California which also found that higher soil organic carbon does not relate to higher wheat yields (Kelley et al., 2024) and with a survey of European farms which found a poor association between soil organic matter and crop yields (Vonk et al., 2020). Future studies could investigate under which conditions of nutrient and water availability lower organic matters can sustain similar to higher yields as high organic matter systems. In addition to higher yields, the cropping systems with S1 also emitted less CO_2_ than S2 systems, which is most likely explained by the enhanced microbial biomass in the S2 soils (ESDAC, 2020). This underlines that definitions of “healthy” cropping systems cannot solely rely on high humus/ soil C content and increased microbial biomass C contents, as these often link to higher microbial activity and therewith accelerated soil CO_2_ emissions, a potentially negative feedback loop to climate change (Hamamoto et al., 2022; Moitinho et al., 2021). Therefore, it is critical to consider the relationship between soil carbon content and nutrient cycling, as these links are fundamental for understanding soil health and its role in biogeochemical cycles (Schröder et al., 2016; Rocci et al., 2024). Overall, the here observed divergence in yields as a function of soil type demonstrates the sensitivity of cropping systems to climate change (Benton, 2020; Hamamoto et al., 2022). To avoid detrimental losses of yield when agricultural systems cross tipping points and to ensure food security worldwide even under climate change, it needs to be carefully evaluate which management practices provide resilience to cropping systems in the long-term (Kornhuber et al. 2023).

### Quality of harvested grains

To prevent malnutrition, it will also be important to determine the technological properties and nutritious value of the harvested grains (Lowe, 2021). In this study, grain nitrogen content was not statistically different between soil types or climates but tended to decrease the further in the future the climate scenario (Table 3). Such a trend of decreasing nutrient levels with increasing temperatures and CO_2_ is often reported in the literature, but our results provide an interesting example of how crops can compensate dilution effects from CO_2_-fertilisation, which merits further study (Liang et al., 2019; Govindasamy et al., 2023). Interestingly, in this study thousand grain weight (TGW), which is an important component of crop yield linked to potential flour yield (Bordes et al., 2008), increased in parallel to the overall yield for the future climate scenarios, but had no consistent relation with soil type (Table 3). This presents a particular challenge for breeding programs, as TGW is closely linked to the stress resilience of wheat plants. Factors like drought and nutrient deficiency can lead to premature ripening and reduced TGW, while also influencing seedling vigour in the next generation (Shahwani et al., 2014). However, developing a soil management plan to target these traits requires more empirical data to establish consistent trends between soil management practices and nutrient redistribution during grain filling, taking the shorter growth cycle in the future climates into account.

### Naturally balancing the nitrogen cycling?

N_2_O is a relevant greenhouse gas with much higher global warming potential than CO_2_ (IPCC, 2014). Agricultural land is one of the main N_2_O sources and as higher temperatures stimulate mineralization and nitrification processes and therewith substrate availability for denitrification, increases in N_2_O emissions are expected in the future (Lamprea et al., 2021). However, depending on local conditions such as soil moisture and N availability, reductions in soil N_2_O emissions are also possible (Muños et al., 2010; Reay et al., 2012). In this study, the DNDC model predicted an overall decrease in soil N_2_O emissions for future climates, with higher N_2_O-emissions in S2 compared to S1, while the difference between S1 and S2 regarding N_2_O-emissions did not change between the years (Fig.3S). As the future years were smoother in the rainfall distribution, the soils were overall drier (Fig.3D,H) and therewith the risk of waterlogging and anaerobic conditions was reduced, limiting denitrification and N₂O production (Venterea, 2007). Moreover, there was no indication of limited plant nitrogen uptake in the future climates (Table 3), which could indicate that the increased soil mineralisation did not lead to nutrient losses in GHGs, but instead nitrogen was taken up by the growing plants. Another mechanism which could have successfully prevented gaseous N-losses could be associated with greater N immobilization belowground, with surplus inorganic NO_3_-N being incorporated into microbial biomass, supporting greater N recycling and retention (Buckeridge et al., 2020; Cao et al., 2021; Pausch et al. 2024). The process of enhanced nutrient immobilisation in microbial biomass could also explain the decreased risk of nitrate leaching predicted for S2 as compared to S1, which has higher microbial biomass, while the sandy texture of S1 likely contributed to its higher susceptibility to nutrient leaching (Gaines & Gaines, 1994). This means that while low organic matter input may be a soil management strategy to reduce GHG-emissions without compromising yield, soils with higher drainage may experience greater nitrate loss through leaching due to the soils’ reduced capacity to retain nutrients effectively. To reduce the environmental impact of high-yielding cropping systems in the future it is therefore essential to identify soil management practices which allow nutrient buffering without necessarily increasing organic matters, like biochar, clay amendments, fertigation or biological nitrification inhibition.

### Stimulating natural plant adaptation to mitigate stress

Plants in the future climatic conditions were more prone to stress because of their changed phenology and a higher potential for disease propagation. For example, leaf area index (LAI) increased significantly under future climate scenarios, which not only indicates increased rates of photosynthesis, but a larger leaf area also allows for greater stomatal conductance, which facilitates more water vapor loss through transpiration and may thus stress plants and increase the water requirements of the cropping system (Zhang et al., 2021). Therewith, plants with larger LAI are more vulnerable to drought spells which can be detrimental for yields (Farooq et al., 2009; Li et al., 2023). Similarly, taller growing plants as observed in the future climates in this experiment are more vulnerable to physical damage from wind and require better rooting systems and stronger cell walls to withstand high wind speeds (Zhao et al., 2022; Jia et al., 2021; Gardiner et al., 2016). In addition, plant health is expected to worsen under climate change when autumns and winters become milder and wetter which favours water logging and the spread of fungal diseases which may ultimately pose severe risks to food production systems (Chakraborty & Newton, 2011). In this study, the pathogenic root fungus *Gaeumannomyces tritici* which causes take-all disease was enhanced in the two future climates, most notably in the 2085 climate, and there particularly in S1, but the pathogenic root infestation did eventually not threaten the yields (Fig. 3L, 3Q). This indicates that the plants could develop successful stress mitigation strategies to combat the pathogen. Two promising factors which could have contributed to the plants’ successful defence are proline and silicon. Proline functions as a potent antioxidant, scavenging reactive oxygen species (ROS) and reducing oxidative damage, which can help protect cellular structures and macromolecules during stress conditions (Hayat et al., 2012). Moreover, exogenous proline application has been shown to improve photosynthetic parameters such as chlorophyll content, stomatal conductance, and PSII efficiency in stressed plants (Sporman et al., 2023) and proline accumulation is associated with increased resistance to various pathogens acting as a signalling molecule triggering defence responses and the production of antimicrobial compounds (Kaur & Asthir, 2015). Silicon on the other hand can contribute to plant health by impregnating cell walls and thus forming a barrier that impedes pathogen penetration, while it also physically strengthens plant stems which could benefit crops that grow taller and are expose to higher wind speeds (Wang et al., 2017). Silicon has also been shown to improve drought tolerance which could become even more important in the future as crops grow with larger leaf area like in this study (Wang et al., 2021) and like proline has been related to oxidative stress mitigation (Kim et al., 2017).

### Implications of phenological advance for farming practices

Plants use various signals to respond to environmental changes, including solar signals which determine the photoperiod, past seasonal experiences like winter chilling, and current conditions such as temperature and moisture (Tang et al. 2016). The advancement in harvest date of approximately two weeks between each climate scenario observed in this study reflects this response and aligns with trends observed in other experiments which report that warmer temperatures accelerate crop development and lead to earlier maturity dates and that in spring photoperiod and winter chilling work together to determine plant growth (Tang et al. 2016; Harkness et al., 2020). The here observed phenological advance in future meteorological conditions has significant implications for agricultural management, for example harvesting winter wheat up to four weeks earlier requires adapting the crop rotation cycle including sowing dates and identifying wheat varieties with shorter maturation cycles. Another observation in this experiment was that plants in 2085 reached heights nearly double those recorded in 2013, which is in line with other studies (Quan et al., 2024). These taller plants may require more physical protection against wind by means of hedges or trees and also imply that harvesting equipment may need to be adapted to the higher positioning of the grains (Miller et al., 2022). Overall, the projected advances in crop maturity and plant height could have profound implications for agricultural practices and as climate conditions continue to evolve, further research will be needed to refine models predicting phenological responses and to develop adaptive strategies for sustainable agricultural practices in changing environments.

### Ecotron constraints

While Ecotrons are unique tools to study agroecosystems under climate change, they are a compromise operating at an intermediate scale between simplistic microcosm experiments and real-world ecosystems, which still cannot fully replicate the complexity of natural environments (Roy et al., 2021; Schmidt et al., 2021). It is therefore vital to cross-validate observations from Ecotron experiments with data from field experiments and to replicate experiments sufficiently. A further challenge to Ecotron experiments is the timescale, as exposing soil and miniature agroecosystems to future meteorological conditions without several generations of adaptive changes occurring implies a very abrupt change in climate conditions. In this experiment for example, soil monoliths sampled in November 2022 had about one month to adapt to the conditions of the simulated November 2085 rather than slowly evolving through 63 years of climate change. However, the experiment with its accelerated evolutionary approach remains realistic, especially for wheat as a plant with a long growth cycle in a crop rotation, because even in current meteorological conditions, year-to-year climatic variability can result in substantial shifts in meteorological conditions between two wheat seasons (Fig. 2). Future experiments could look at how the accumulation of several more extreme seasons affects cropping systems and how the occurrence of individual more extreme events within one growing season impacts crop performance.

## Conclusion

This study provides new insights into the complex interplay between climate change, soil management practices, and winter wheat performance. The observed phenological advances, increased yields in low organic matter soils (S1) as compared to high organic matter soil (S2), and natural stress adaptation mechanisms such as proline and silicon accumulation highlight the capacity of crops to respond to future climate conditions. However, additional research is needed to better understand why in some soil conditions (like S2 in this study) the CO_2_-fertilization effect remains limited, and yields decrease under future climates. This may be linked to higher nutrient immobilisation in high organic matter systems compared to low organic matter systems, which could also explain the lower risk of nitrate leaching in S2. To ensure sustainable performance of future cropping systems it would therefore be key to further develop management practices that allow fertiliser application and nutrient cycling without necessarily increasing organic matters because of the enhanced CO_2_-emissions and nutrient immobilisation. Fertigation or biological nitrification inhibition could be tested under future meteorological conditions to assess their potential to optimise nutrient availability and use efficiency in agroecosystems. This study further highlights the need to investigate links between altered soil processes and plant diseases under future climates, where shorter cropping cycles may provide an opportunity to break disease cycles and restore soil nutrient stocks with inter-cropping.

## Supporting information captions

**SM1**: Soil management history

**SM2**: Monolith sampling

**SM3**: Fertilization during Ecotron experiment

**SM4**: Details on methods of empirical and modelled parameter to quantify agronomic performance and environmental impact

**SM5**: DNDC model evaluation

**SM6**: Loadings for the main vector of each modality (Year.Soil)

## Supplementary material for

**SM1**: Soil management history

**SM2**: Monolith sampling

**SM3**: Fertilization during Ecotron experiment

**SM5**: DNDC model evaluation

**SM6**: Loadings for the main vector of each modality (Year.Soil)

## SM1: Detailed soil management

**Soil type 1 (S1) 50°38’35.1474”N, 4°37’22.0123”E**

| Date | Crop / intervention | Dose | Comments/Details |
| --- | --- | --- | --- |
| 2015 | Wheat |  |  |
| 2016 | Chicories |  |  |
| 2017 | Wheat |  |  |
| 07/08/17 | 1 pass of harrowing |  |  |
| 24/08/17 | Incorporation of Haspargit® and scums |  | Haspargit®, low-chlorine potassium fertilizer, contains sulfur and calcium |
| 05/03/18 | 350 kg/ha of Sulfonitrate N26+ 31 SO3(commercial name) | 350 kg/h |  |
| 06/04/18 | 350kg/ha Nitrogen (N27 %) | 350 kg/h |  |
| 15/04/18 | CARYX® | 1,25 L/ha | Growth Regulator + Fungicide |
| 04/05/18 | Tangus Gold | 0,5 kg/ha | Fungicide |
| 14/07/18 | Canola harvest |  |  |
| 21/08/18 | Harrowing |  |  |
| 03/10/18 | Tillage |  |  |
| 03/10/18 | Plantation escourgeon multi |  |  |
| 06/11/18 | HEROLD® | 0,5 L /ha | HEROLD®, contact and residual broad-spectrum herbicide based on the leading grass-weed active ingredient flufenacet, for pre- and post-crop emergence application |
| 06/11/18 | Chlortoluron | 1,2 L/ha | Herbicide |
| 06/11/18 | Patriot protect | 0,32 L/ha | Patriot® Selective Herbicide |
| 26/02/19 | Nitrogen N27 % | 250 kg/ha |  |
| 28/03/19 | KANTIK® | 1 L/ha | Fungicide |
| 28/03/19 | MEDAX® TOP | 1,2 L/ha | MEDAX® TOP is a growth regulator for cereals |
| 28/03/19 | MAGNESIE | 2,5 kg/ ha |  |
| 02/04/19 | SULFAZOTE 22%N (28,6 U) | 200 L/ha |  |
| 20/04/19 | TERPAL® | 1,2L /ha | Terpal® shortens the internodes and strengthens the stem wall, treated cereal crops become more resistant to lodging |
| 20/04/19 | INPUT | 0,6L /ha |  |
| 02/05/19 | VELOGY ERA | 0,7L / ha | FUNGICIDE |
| 02/05/19 | BRAVO | 1,2L /ha | FUNGICIDE |
| 02/05/19 | MAGNESIE | 2,5KG /ha |  |
| 02/05/19 | Patriot protect | 0,2L /ha | Patriot® Selective Herbicide |
| 08/07/19 | Harvest escourgeon multi |  |  |
| 24/08/19 | Glyfall plus | 6L/ha | Total herbicide |
| 02/09/19 | Harrowing |  |  |
| 09/09/19 | Harrowing |  |  |
| 11/09/19 | Sowing seeds (radish, clover, phacelia) | 12kg/ha |  |
| 2021 | Sugar beet/wheat culture |  | Variety BTS 4860 + KWS Tessilia + Caprianna kws + Bts 3480 (6 boxes of each) |
| 03/04/20 | Sugar beet sowing |  |  |
| 21/08/20 | Fertilizer 0-5-16+15 CaO+2 MgO+12 EB | 1113 kg/ha |  |
| 02/09/20 | Harrowing |  |  |
| 07/09/20 | Harrowing |  |  |
| 07/09/20 | Liquid nitrogen | 120 l/ha |  |
| 07/09/20 | Green manure seeding 15kg/ha (base 5kg + early 10kg) brand Lemken |  |  |
| 31/10/20 | Fast plow |  |  |
| 06/12/20 | Tillage |  |  |
| 23/03/21 | 7 So3 | 800 kg/ha | Fertilizer |
| 23/03/21 | Nitrogen N27 | 150 kg/ha |  |
| 23/03/21 | 7 So3 | 800 kg/ha | Fertilizer |
| 26/03/21 | Liquide nitrogen | 350 L/ha |  |
| 02/04/21 | Soil preparation |  |  |
| 04/04/21 | Beetix | 1,75 L/kg | Herbicide |
| 04/04/21 | Centium® | 0,05 L/kg | Herbicide |
| 01/05/21 | Dianal | 0,9 L/ha |  |
| 01/05/21 | Ethomat | 0.3 | Herbicide to control annual dicot weeds and annual grass weeds in sugar beet and fodder beet. |
| 01/05/21 | Safari® | 15 gr/ha | Safari® herbicide for post-emergence control of broadleaf weeds in sugar beets, red beets, chicory, and endive |
| 01/05/21 | Goltix® Queen | 1 L/ha | Goltix® Queen is a combination of the active ingredient metamitron, and an addition of the active ingredient quinmerac, long-lasting control on regrowth, effectively controlling a broad spectrum of weeds |
| 01/05/21 | Vegetop | 0,5 L/ha | Additive to enhance herbicide performance. |
| 01/05/21 | Magnesia | 2,3kg /ha |  |
| 07/05/21 | Dianal | 0,9 L/ha |  |
| 07/05/21 | Ethomat | 0.3 | Herbicide to control annual dicot weeds and annual grass weeds in sugar beet and fodder beet. |
| 07/05/21 | Goltix® Queen | 1 L/ha | Goltix® Queen is a combination of the active ingredient metamitron, and an addition of the active ingredient quinmerac, long-lasting control on regrowth, effectively controlling a broad spectrum of weeds |
| 07/05/21 | Vegetop | 0,5 L/ha | Additive to enhance herbicide performance. |
| 07/05/21 | Magnesia | 2,3kg /ha |  |
| 11/05/21 | Hoeing |  |  |
| 29/05/21 | Dianal | 0,9 L/ha |  |
| 29/05/21 | Ethomat | 0.3 | The product is intended to be used as herbicide to control annual dicot weeds and annual grass weeds in sugar beet and fodder beet. |
| 29/05/21 | Beetix | 0,6 L/ha L/kg | Herbicide |
| 29/05/21 | Tamaris | 0,6 L/ha |  |
| 29/05/21 | Lenazar | 150 gr/ha | Selective herbicide against annual dicotyledonous weeds |
| 07/05/21 | Vegetop | 0,5 L/ha | Additive to enhance herbicide performance. |
| 15/06/21 | Centium® | 0,05 L/kg | Herbicide |
| 15/06/21 | Frontier® Optima | 0,5 L/ha | Frontier Optima is a soil herbicide primarily absorbed through the coleoptile of grasses as the seedling penetrates the upper layer of the soil. |
| 15/06/21 | Boron | 3 L/ha | Fertilizier |
| 15/06/21 | Magnesia | 2,3kg /ha | Fertilizier |
| 05/08/21 | Agora | 0,35l /ha | Fungicide |
| 05/08/21 | Magnesia | 5,9kg /ha |  |
| 01/09/21 | Bicanta | 1 L/ha | Bicanta: product to combat diseases in sugar beet cultivation, combination of two active ingredients 125 g/l of difenoconazole and 125 g/l of azoxystrobin, effective control against all major foliar diseases, including Cercospora, Stemphylium, downy mildew, Ramularia, and rust |
| 01/09/21 | Magnesia | 5 kg/ha |  |
| 16/10/21 | Harvest |  |  |
| 15/11/21 | Harvest |  |  |
| 2022 | Wheat |  |  |
| 18/10/21 | Sowing wheat | 153 kg/ha | Variety Campesino PMG:44 gr |
| 06/11/21 | Thin sowing wheat |  |  |
| 01-03-22 | 18 SO3 | 260 kg/ha |  |
| 15-03-22 | Sulfazote 22% | 200 l/ha |  |
| 23-03-22 | Osmose | 0,2 l/ha |  |
| 23-03-22 | SAVVY® | 20 gr/ha | Herbicide, 200g/kg of Metsulfuron-methyl |
| 23-03-22 | Sigma® Maxx | 1,1 L/ha | Selective post-emergence herbicide in winter wheat, spring wheat, rye, triticale, and spelt against annual grasses and dicotyledons |
| 23-03-22 | Magnesia | 2 kg/ha |  |
| 05-04-22 | Nitrogene N27 % | 194 kg/ha | Fertilizer |
| 18-04-22 | FERTILEADER® Trio |  | Improves plant resistance to stress, stimulates metabolism, improves plant photosynthesis, contains patented Seactiv® complex (trademark of Timak Agro) |
| 18-04-22 | Magnesia | 2,3 kg/ha |  |
| 18-04-22 | Osmose | 0,2 l/ha |  |
| 18-04-22 | Chlormequat | 1 L/ha | Growth regulator, can control the vegetative growth of plants (i.e., root and leaf growth), promote the reproductive growth of plants (i.e., flower and fruit growth), and enhance the plant's fruit-setting rate. |
| 27-04-22 | Tebecur | 2,7 kg/ha | All-round fungicide with efficacy over a period of several weeks, controlling numerous pathogens in various crops, effective on i.a. Fusarium, Alternaria, Septoria nodorum, rusts, Sclerotinia, powdery mildew, leaf spots |
| 27-04-22 | Osmose | 0,2 l/ha |  |
| 27-04-22 | Magnesia | 2,7 kg/ha |  |
| 15-05-22 | Nitrogen N38 | 31 L/ha | Foliar fertilizer |
| 18-05-22 | Osmose | 0,2 l/ha |  |
| 18-05-22 | Magnesia | 3,6 kg/ha |  |
| 18-05-22 | REVYTREX® | 1,1 L/ha | New cereal fungicide based on Revysol® |
| 18-05-22 | COMET® | 0,4 L/ha | Preventive-action fungicide designed to combat foliar diseases in various cereals (eg fight dwarf rust) |
| 29/07/22 | Harvest wheat |  |  |
| 05/08/22 | Harrowing |  |  |
| 05-08-22 | KALCIPHOS P-K (S) | (0-5-15)<br>1000<br>kg/ha |  |
| 19/08/22 | Cattle manure | 15t/ha |  |
| 24/09/22 | Green fertilizer | 11,6 kg/ha |  |
| 24/09/22 | Fast plowing |  |  |

**Soil type 2 (S2) 50°39’12.8668”N, 4°38’10.7664”E**

**Soil type 2 (S2) 50°39'12.8668"N, 4°38'10.7664"E**
| Date | Crop / intervention | Dose | Comments/Details |
| --- | --- | --- | --- |
| 2017 | Beet culture |  |  |
| 2018 | Wheat culture |  |  |
| 2018 | Cover crop after harvest (oats, mustard, radish, clover, sunflower, peas). |  |  |
| 2019 | Potato culture |  |  |
| 2019-2020 | wheat |  |  |
| 20/10/19 | Mixed vegetation seeding | 145 kg/ha |  |
| 02/03/20 | Nitrogen N39 | 175 L/kg |  |
| 01/02/20 | TMS | 80 kg/ha | TMS, mineral CE-fertilizer, regulates microbial flora and organic matter evolution, improves chemical, physical and biological soil fertility. |
| 16/03/20 | Sigma® Flex | 140 g/ha | Herbicide (not sure about the unit looks like g/ha) |
| 16/03/20 | Brodway | 150 g/ha | Herbicide (not sure about the unit looks like g/ha) |
| 16/03/20 | Oil | 1l/ha |  |
| 06/04/20 | Cycocel |  | Cycocel is a plant growth regulator |
| 18/04/20 | Nitrogen N39 | 170 L/ha |  |
| 18/04/20 | Humic and fulvic acids | 1 L/ha | Fertilizer |
| 16/05/20 | Tebuzifo | 04 L/ha |  |
| 16/05/20 | TMF | 1L/ha | Fertilizer |
| 16/05/20 | MgSo4 | 1,8 kg/ha | Fertilizer |
| 20/05/20 | Nitrogen N 27 | 200 kg/ha | Fertilizer |
| 02/06/20 | MgSo4 | 1,8 kg/ha | Fertilizer |
| 02/06/20 | TMF | 1L/ha | Fertilizer |
| 02/06/20 | Opustear | 0,35 l/ha |  |
| 02/06/20 | Sportitck | 0,36 l/ha |  |
| 02/06/20 | Kestrel | 0,36 l/ha |  |
| 20/08/20 | Cow manure | 12000L | Fertilizer |
|  | Cover crop seeding (phacelia, oats, radish, clover) |  |  |
|  | Grazing of cover crops, CRAW Janart trials |  |  |
| 08/03/21 | Glypho | 1L/ha | Herbicide |
| 23/02/21 | TMS | 80 kg/ha | Fertilizer |
| 03/03/21 | Unreadable | 300 kg |  |
| 18/03/21 | Nitrogen | 100L/ha | Fertilizer |
| 18/03/21 | Humactiv | 3,5 L/ha | Root stimulant |
| 11/05/21 | Flax |  |  |
| 11/05/21 | Kaei |  |  |
| 11/05/21 | TMS | 1 L/ha |  |
| 11/05/21 | MgSo4 | 3kg/ha |  |
| 11/06/21 | Rudis® | 1 L/ha | Fungicide |
| 11/06/21 | TMF | 0,4 L/ha | Fertilizer |
| 11/06/21 | éthéplum | 4 L/ha |  |
| 15/06/21 | TOPREX | 0,3 L/ha | TOPREX, co-formulation of difenoconazole and paclobutrazol, use as plant growth regulator, also as a fungicide against phoma and light leaf spot in winter oilseed rape |
| 15/06/21 | Flax |  |  |
| 16/10/21 | Wheat Sowing |  |  |
| 05/11/21 | Liberator + Flufenacet | 0,58 L/ha | Herbicides |
| 05/11/21 | Mateno® Duo | 0,35 L/ha | Herbicides |
| 26/02/22 | TMS | 80 kg/ha | Fertilizer |
| 14/03/22 | Nitrogen N39 | 200 L/ha | Fertilizer |
| 05/04/22 | Nitrogen N39 | 175 L/ha | Fertilizer |
| 13/04/22 | MgSO4 | 3,5 kg/ha | Fertilizer |
| 13/04/22 | TMF | 1 L/ha | Fertilizer |
| 13/04/22 | Cycocel | 1L /ha | Cycocel is a plant growth regulator |
| 25/04/22 | Nitrogen N39 | 125 L/ha | Fertilizer |
| 22/05/22 | Kestrel | 1 L/ha |  |
| 22/05/22 | MGSO4 | 3kg/ha | Fertilizer |
| 22/05/22 | TMF | 1L/ha | Fertilizer |
| 01/06/22 | Tebuzip ?? Maybe Tebuconazole | 0,7 L/ha | ?? |
| 01/06/22 | TMF | 1L/ha | Fertilizer |
| 01/06/22 | Prinus | 0,1 ?? |  |
|  | Wheat | 108 Qx/ha |  |
| 15/08/22 | Horse manure | 15 T/ha | Fertilizer |
| 17/09/22 | Mixture Scam Selfie |  | Cover crop |

SM2: Soil monolith sampling

**SM2: Soil monolith sampling**

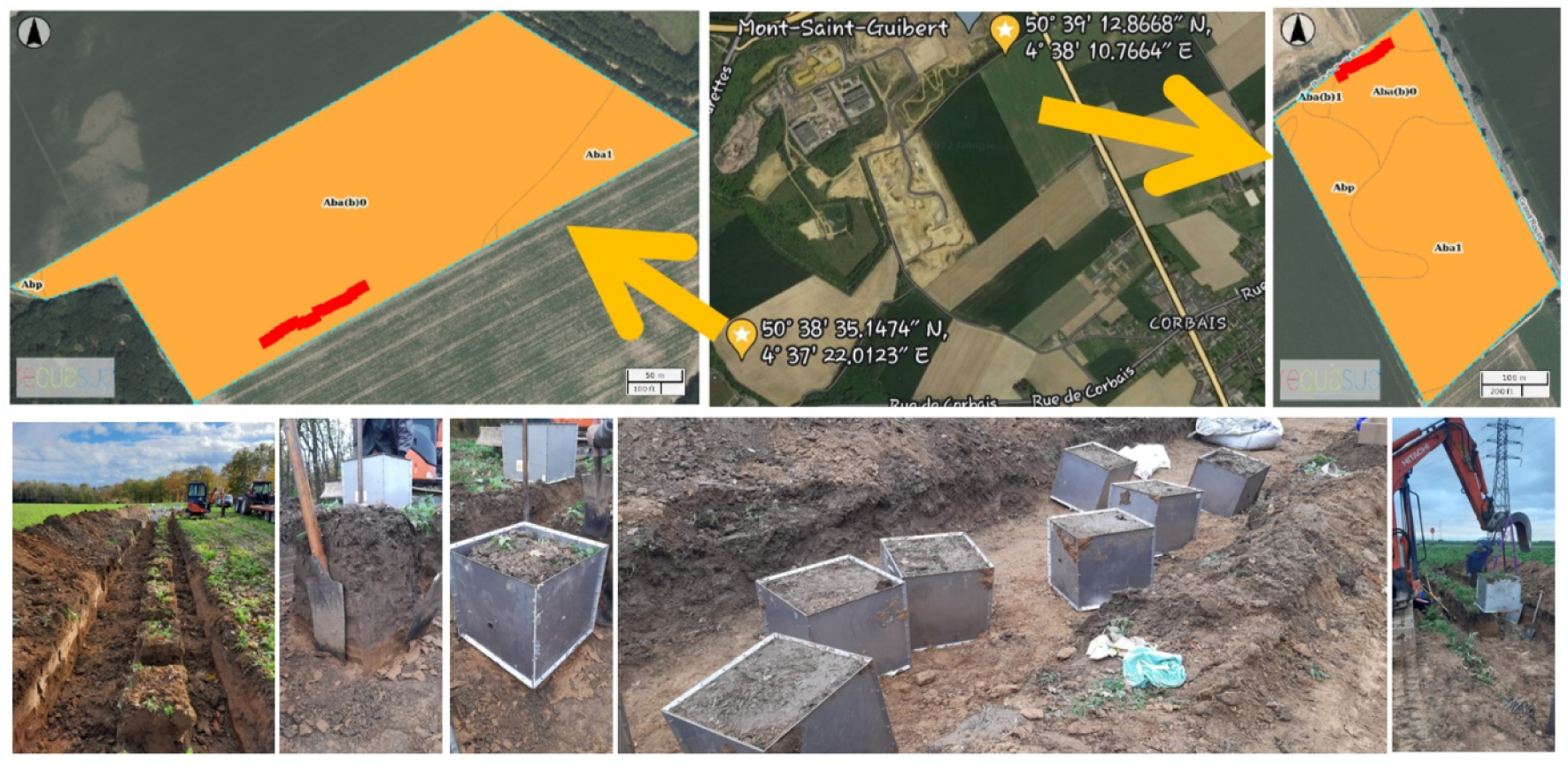

**SM3: Fertilisation in BIOFAIR Ecotron trial**

The total required amount of nitrogen fertiliser was split into three doses, each of which was applied according to plant growth stage in respective climate. The recommendation was made for each soil type in each CER and based on the guidelines of the Wallon Centre of Agronomic Research (CRA-W) as described in Le Livre Blanc: La fertilisation azotée. The recommendation consists in calculating a forecast N balance for the crop, taking into account the soil nitrogen supplies and the estimated needs of the intended crop. It makes it possible to assess the adequate quantity of fertilizer to bring. Soil nitrogen supplies are determined based on soil characteristics (humus content, remaining mineral nitrogen at the end of winter, mineralization of organic nitrogen in the soil), the phytotechnical history of the plot (crop residues, previous crop) and mineral and previous organic nitrogen fertilizer inputs by the farmer.

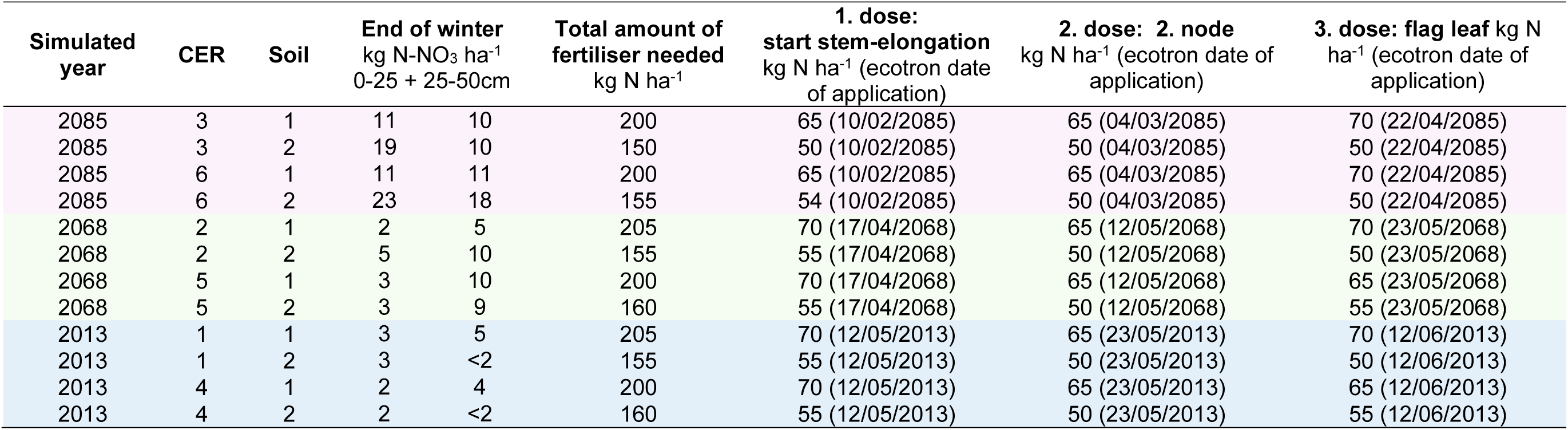

**SM4: Empirical and modelled parameter to quantify agronomic performance and environmental impact**

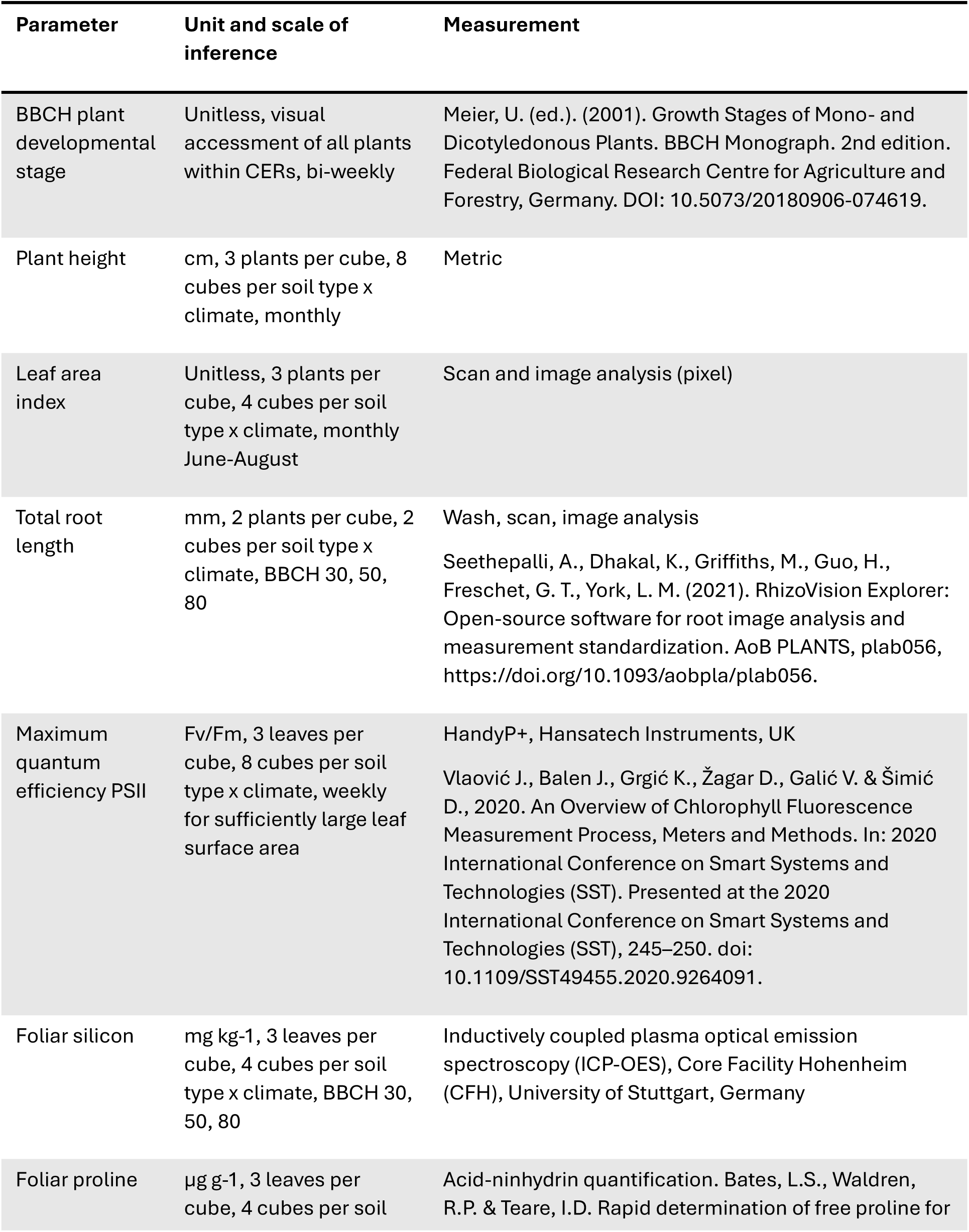
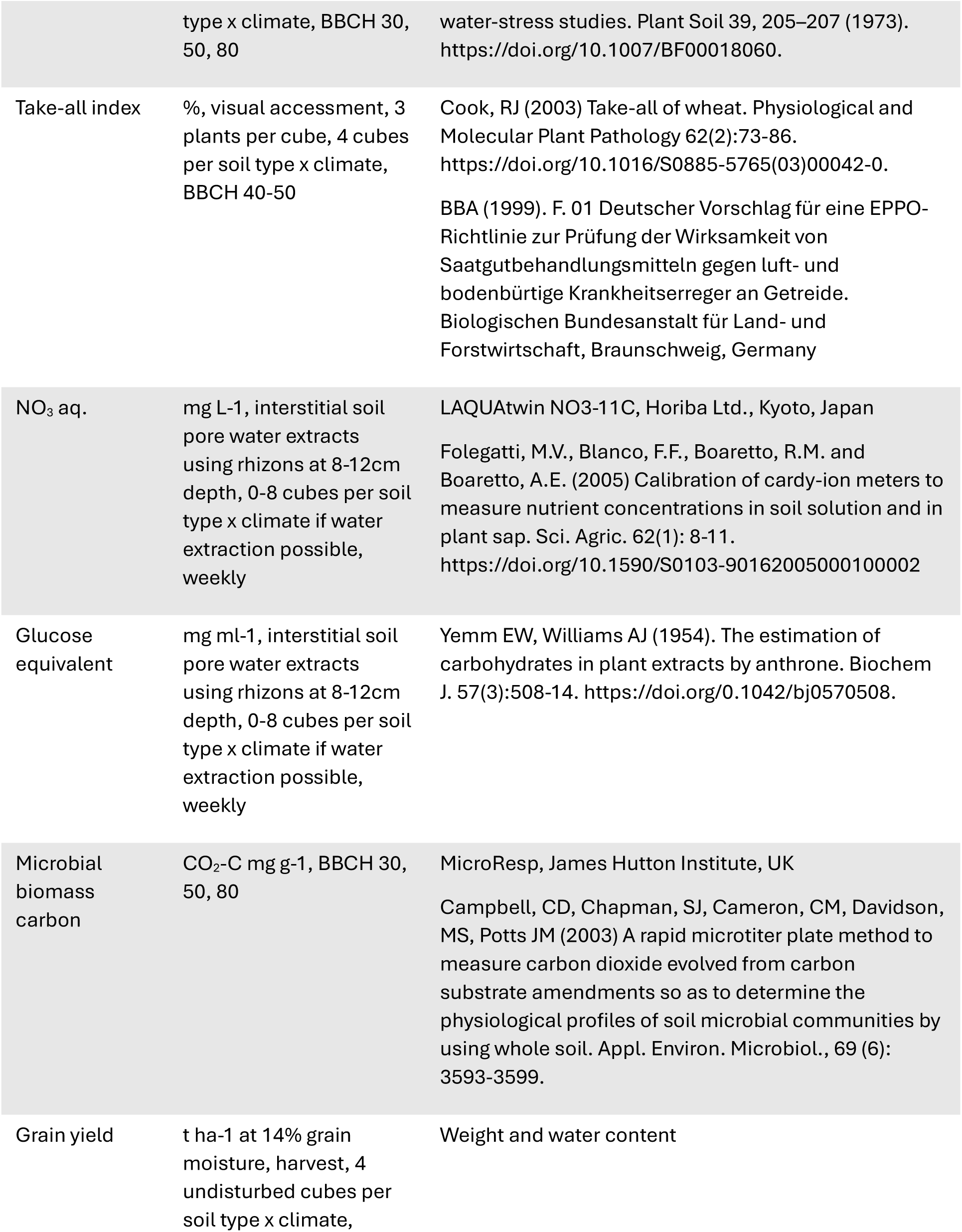
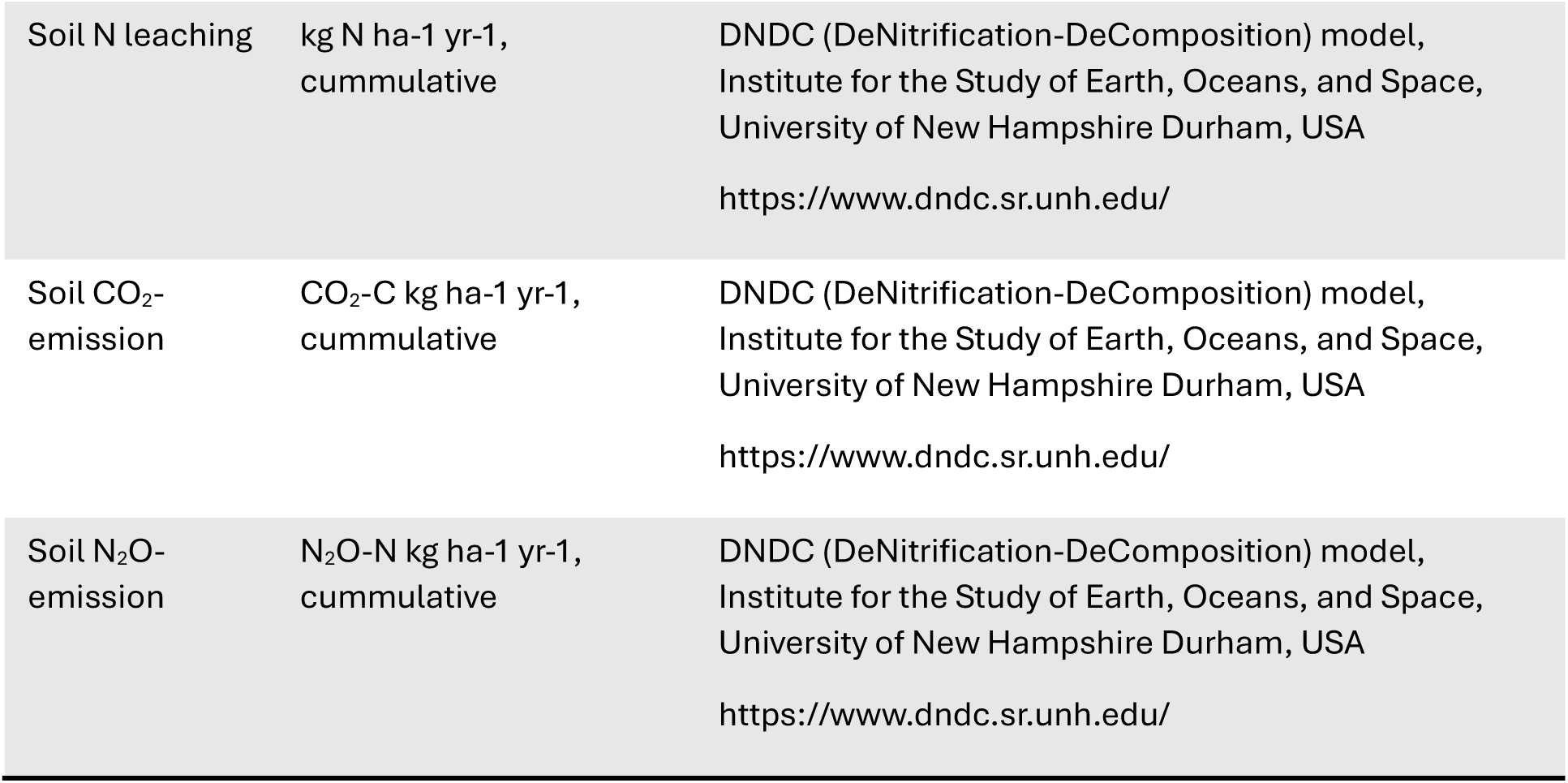

**SM5: DNDC model evaluation against measured data**

To evaluate the accuracy of the modelled parameters from the DNDC model, the results were compared against measured values using the coefficient of determination (R^2^). The correlation was excellent for the DNDC estimate of CO_2_-emission and the basal soil microbial respiration accessed with the MicroResp kit (R^2^=0.98). While no empirical data directly on denitrification was gathered during the Ecotron experiment, the DNDC estimate of soil N_2_0-emission correlated well with the metabolic quotient (qCO_2_), probably reflecting that microbes that actively denitrify (under anaerobic conditions) also engage in respiration, leading to CO_2_ emissions (R^2^=0.6). The correlation between the measured nitrate in aqueous solution and the predicted soil N-leaching was good (R^2^=0.71) if 2013.S1 was not taken into account. Similarly, a moderate correlation (R^2^=0.53) was observed for the root exudation predicted by the DNDC model and the measured values of glucose equivalent, if 2013.S1 was excluded (R^2^=0.53). The data distribution suggests that the NO_3_ in aqueous solution in 2013.S1 was actually higher than what was measured, which could be related to the fast turnover time of NO_3_ in soil where it is quickly taken up by plants and microbes. For root exudation, in DNDC root exudates are represented as a portion of the root-derived carbon where the model considers both root growth and root turnover, while the empirical measurement performed here only gives an estimate of low-molecular weight sugars measured as glucose equivalents which may explain discrepancies. The data distribution suggests that both the DNDC model and the measurements underestimated root exudation in 2013.S1, which similarly to NO_3_ could be due to the high reactivity of these molecules in the soil. Overall, the modelled and measured parameter were well aligned for gaseous emissions of CO_2_ and N_2_O (R^2^=0.98, R^2^=0.6), and moderately well aligned for nitrate and glucose equivalent (R^2^=0.71, R^2^=0.53) with 2013.S1 not fitting the distribution, possibly due to the technical limitations of the empirical measurements.

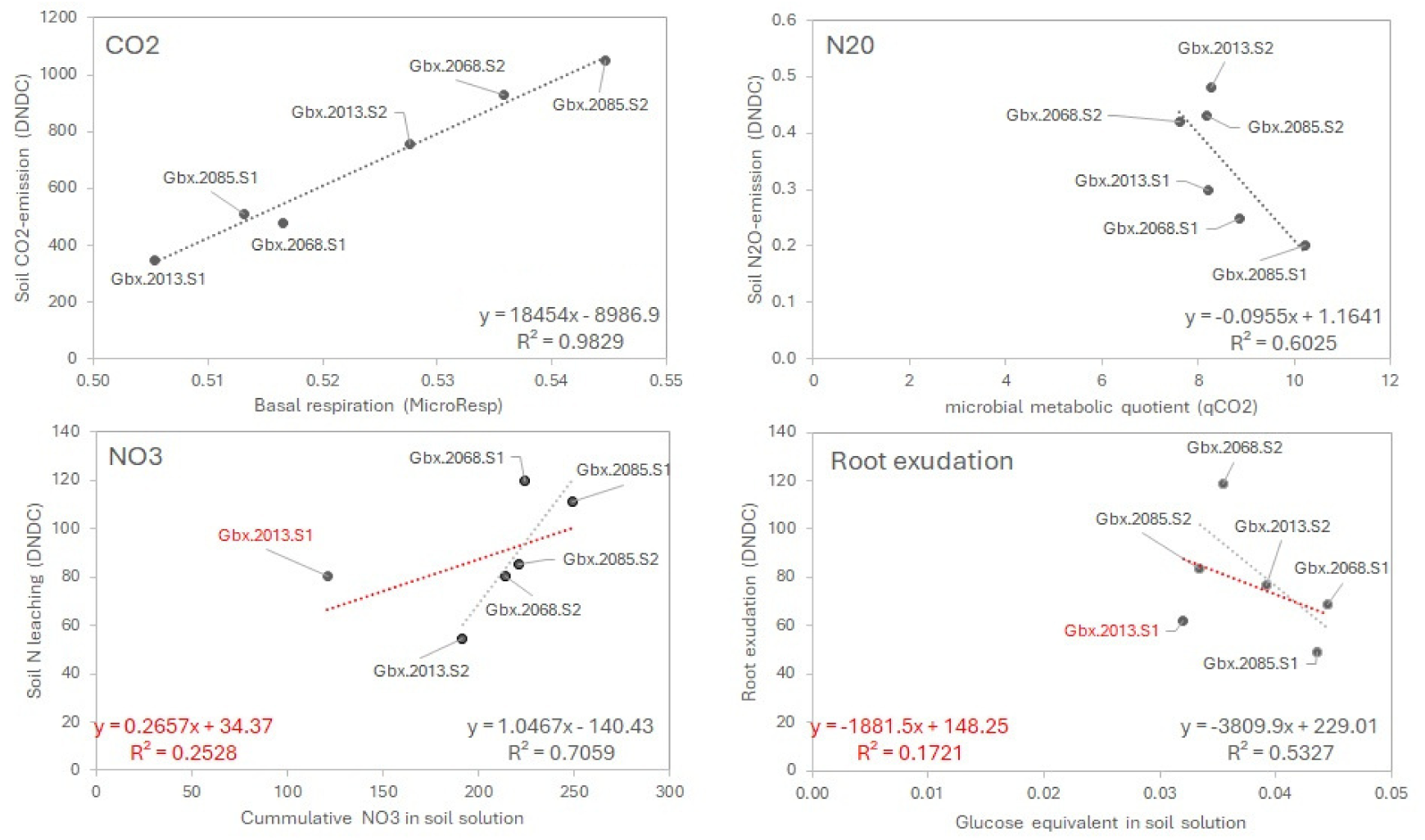

**SM6: Loadings for the main vector of each modality (Year.Soil)**

|  | 2013.S1 | 2013.S2 | 2068.S1 | 2068.S2 | 2085.S1 | 2085.S2 |
| --- | --- | --- | --- | --- | --- | --- |
| Basal.CO2.02.05 |  |  |  | -0.8399 |  | 1.55331 |
| Basal.CO2.30.05 | -0.7269 | -0.8814 |  | 1.23257 |  |  |
| Basal.CO2.03.07 |  |  |  | -0.8972 | 1.25774 |  |
| Basal.CO2.07.08 |  |  | -0.0394 |  |  |  |
| Foliar.proline_BBCH80 |  |  | 0.14672 |  |  |  |
| Foliar.silicon_BBCH50 |  |  |  |  |  | 1.36197 |
| Glucose.equivalent.27.02 | 0.82706 |  | -0.1687 |  |  |  |
| Glucose.equivalent.14.03 | -0.6393 |  |  |  |  |  |
| Glucose.equivalent.02.05 | 0.67934 |  |  | -0.7745 |  |  |
| Glucose.equivalent.21.05 |  | -0.896 |  |  |  |  |
| Glucose.equivalent.03.07 |  |  | -0.0938 |  |  |  |
| Glucose.equivalent.19.07 | -0.7882 | -1.0001 |  |  |  |  |
| Glucose.equivalent.25.08 |  |  | -0.1407 |  |  |  |
| Leaf.area.index.March | -0.864 | -0.8922 |  |  |  | 1.31855 |
| Leaf.area.index.April |  | -1.0344 | 0.09927 |  |  | 1.53179 |
| Leaf.area.index.May |  | -1.141 |  |  |  |  |
| Leaf.area.index.June |  | -1.1419 |  |  |  |  |
| Maximum.quantum.efficiency.PSII.23.03 |  |  |  | -0.7742 | 1.17837 | 1.12584 |
| Maximum.quantum.efficiency.PSII.07.04 |  |  |  | 0.82972 |  | -1.2996 |
| Maximum.quantum.efficiency.PSII.19.04 |  |  | -0.1273 |  |  |  |
| Maximum.quantum.efficiency.PSII.10.05 |  |  |  | -0.9076 |  |  |
| Maximum.quantum.efficiency.PSII.08.06 |  | -0.9777 |  |  |  |  |
| Maximum.quantum.efficiency.PSII.21.06 | -0.8253 | -1.2397 |  |  |  |  |
| Maximum.quantum.efficiency.PSII.05.07 |  |  |  | 0.89958 |  |  |
| Microbial.biomass.carbon_BBCH30 |  |  | -0.1362 |  |  |  |
| Microbial.biomass.carbon_BBCH50 |  |  | 0.03226 | 0.72059 |  |  |
| NO3.aq.14.03 |  |  |  |  | -1.3856 |  |
| NO3.aq.15.05 | 0.75471 |  |  | -0.8281 |  |  |
| NO3.aq.12.06 |  |  |  |  | 1.53893 |  |
| NO3.aq.03.07 |  | -0.9794 |  |  |  |  |
| NO3.aq.19.07 | -0.7774 |  |  |  |  |  |
| NO3.aq.07.08 |  |  |  |  | 1.16468 |  |
| Plant.height.March |  |  |  |  | 1.4301 | 1.14948 |
| Plant.height.April |  |  |  |  | 1.43003 | 1.14928 |
| Plant.height.May |  |  |  |  | 1.43006 | 1.14936 |
| Plant.height.June |  |  |  |  | 1.43012 | 1.14976 |
| Take.all.index | -0.722 |  |  |  |  |  |
| Total.root.length_BBCH30 |  |  | -0.0929 |  | 1.13926 |  |

## Notes

### Competing Interest Statement

The authors have declared no competing interest.

### Summary of Updates

Changed title and abstract to clarify soil organic matter management strategies

